# Micro-scale fluid behavior during cryo-EM sample blotting

**DOI:** 10.1101/791285

**Authors:** M. Armstrong, B-G. Han, S. Gomez, J. Turner, D. A. Fletcher, R. M. Glaeser

## Abstract

Blotting has been the standard technique for preparing aqueous samples for single-particle electron cryo-microscopy (cryo-EM) for over three decades. This technique removes excess solution from a TEM grid by pressing absorbent filter paper against the specimen prior to vitrification. However, this standard technique produces vitreous ice with inconsistent thickness from specimen to specimen and from region to region within the same specimen, the reasons for which are not understood. Here, high-speed interference-contrast microscopy is used to demonstrate that the irregular pattern of fibers in the filter paper imposes tortuous, highly variable boundaries during removal of excess liquid from a flat, hydrophilic surface. As a result, aqueous films of nonuniform thickness are formed while the filter paper is pressed against the substrate. This pattern of nonuniform liquid thickness changes again after the filter paper is pulled away, but the thickness still does not become completely uniform. We suggest that similar topological features of the liquid film are produced during the standard technique used to blot EM grids and that these manifest in nonuniform ice after vitrification. These observations suggest that alternative thinning techniques, which do not rely on direct contact between the filter paper and the grid, may result in more repeatable and uniform sample thicknesses.

**STATEMENT OF SIGNIFICANCE:** Multiple imaging techniques are used to observe dynamic, micro-scale events as excess water is removed from a substrate by blotting with filter paper. As a result, new insight is gained about why the thickness values of remaining sample material are so variable across a single EM grid, as well as from one grid to the next. In addition, quantitative estimates are made of the shear forces to which macromolecular complexes can be exposed during blotting. The fact that sample thicknesses and flow rates are seen to be inherently under poor control during blotting suggests that other methods of removing excess water may be better suited for consistently achieving large sample areas that are suitable for use in electron cryo-microscopy.

## INTRODUCTION

The prevailing method of making thin specimens for electron cryo-microscopy (cryo-EM) involves blotting excess sample by pressing filter paper against one or both sides of the specimen grid (EM grid), and then rapidly freezing (vitrifying) the resulting thin film of liquid that remains on the grid (1). Although this has been the established method of preparing cryo-EM grids for over 30 years, there still is little theoretical understanding or direct observation of what occurs during this process.

A recognized shortcoming of this method is that the thickness of the sample is much too variable from one grid to the next, and even within one grid. As a result, one might have to prepare and screen a number of grids before one is found for which the ice thickness is suitable for high-resolution data collection over an acceptable fraction of the area. This lack of reliability has led recent efforts to investigate entirely new ways to make uniformly thin films, including spraying small-volume droplets of samples onto continuous carbon films (2, 3) or onto self-wicking grids (4, 5), and dip-pen or equivalent writing of thin liquid films onto EM grids (6, 7). However, more information is needed about how the standard method fails in order to guide development of alternative approaches. Here we seek to improve our understanding of the microscale-fluid behavior that occurs when filter paper is pressed directly against the face of a 3 mm hydrophilic disc, either an EM grid or a glass coverslip that serves as a proxy for an EM grid.

To this end, we employ a variety of electron-microscope and light-microscope techniques to investigate what happens during blotting. We first compared scanning cryo-EM (cryo-SEM) images to low-magnification transmission EM (TEM) images and observed that variable ice thicknesses occur on both the front (i.e. carbon-film) and back sides of EM grids. Then, using confocal light microscopy, we confirmed that a relatively thick, continuous film of water exists between the holey carbon film on an EM grid and the wet filter paper, provided that there is an excess of sample. Finally, we captured the dynamics of air-finger intrusion between the substrate and filter paper, using high-speed reflection interference-contrast microscopy and a hydrophilic 3-mm glass coverslip as a proxy for an EM grid. We observed that fingers of air rapidly intrude into the previously lubricated gap shortly after blotting begins, when typical volumes of aqueous solution are used.

Our light-microscopy experiments make it clear that randomly distributed points of contact between fibers in the filter paper and the hydrophilic substrate create complex boundaries that govern the removal of the sample. These boundaries determine, as is indicated in Figure 1C, how much water remains on the grid and where, once adsorption into the filter paper is completed (Figure 1). The irreproducibility of these boundary conditions, and the limited extent to which they can be controlled, suggests that progress towards achieving uniform sample thicknesses will require designing alternative techniques to blotting with filter paper.

**Figure 1.**
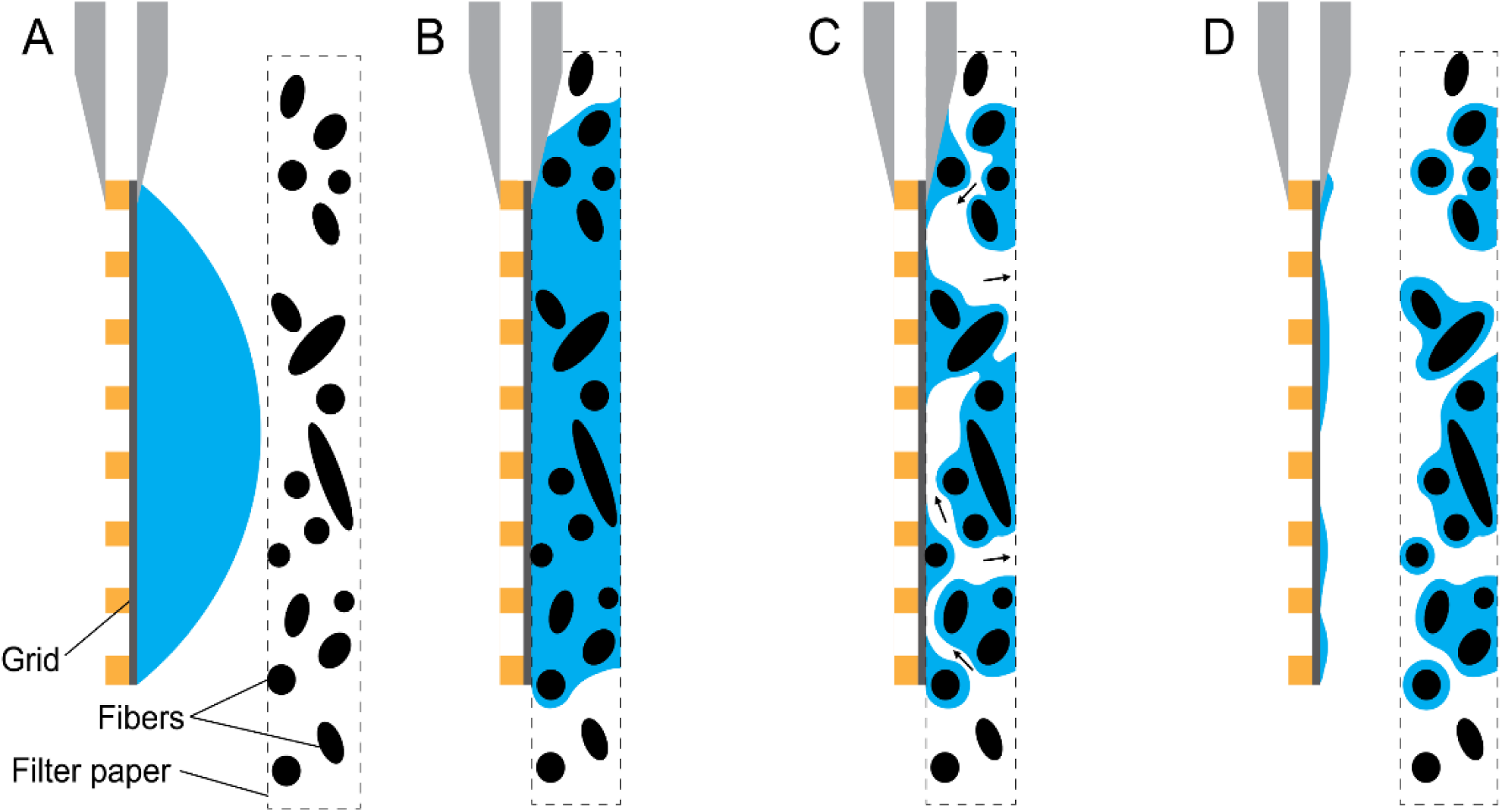
Cartoon summarizing the observed mechanism of sample thinning. A second filter paper, if any, along with any liquid on the back side of the grid is not shown. See the Discussion section for remarks about the extent to which various commercial and home-built implementations may differ from that used here. (A) Before contact with the filter paper, the aqueous sample forms a spherical cap, covering the front side of the grid. (B) When the filter paper is pressed against the grid, a lubrication layer initially remains continuous over the front surface of the grid, with varying distances between the carbon film and the nearest fibers. (C) While the filter paper is still pressed against the grid, capillary forces pull the aqueous sample deeper into to the filter paper, causing fingers of air to propagate into the gap from the edges of the grid, and leaving regions of varying water film thickness across the surface of the grid. (D) As the filter paper is pulled away, the air-water surface collapses locally around points where it previously was constrained by a fiber, but dewetted areas prevent the surface from returning to a meniscus covering the entire area of the grid.

## METHODS

### Evaluation of surface distribution and total thickness distribution of vitreous ice

Low-magnification transmission electron microscope (TEM) images showing the distribution of vitreous ice were obtained with grids prepared in the course of ongoing work to develop streptavidin affinity grids (8). While that meant that the holes in the holey-carbon substrate were spanned by a continuous support film, the results obtained after blotting were characteristic of what is otherwise observed when there is no film over the holes.

A Vitrobot Mark IV (ThermoFisher Scientific, Hillsboro, OR) instrument was used to blot and plunge grids. Blotting was typically done for 5 seconds at a force setting of “10”, a relative humidity of 100% and a temperature of 20 °C, followed by plunging into liquid ethane with zero delay. Varying the blot time or the force setting had no systematic effect on the vitreous ice distribution as reflected by the projected ice-thickness distribution seen in the “grid atlas” image, which was collected with the Leginon software (9).

Scanning electron microscope (SEM) images of selected grids were subsequently obtained with a FEI Strata 235 dual-beam microscope (ThermoFisher Scientific, Hillsboro, OR) equipped with a Quorum 3010 Cryogenic-Transfer (Laughton, UK) system. A homemade device was used to hold previously frozen, standard EM grids on the sample stub. Evidence of crystallization of the previously vitreous ice was seen, indicating that warming to above the glass-to-crystal phase-transition temperature occurred during transfer into the cold stage. Nevertheless, crystallization of the previously vitrified sample did not affect our ability to compare ice distributions on the front and back faces of grids and to compare these distributions to those seen in projection.

### Measurement of the thickness of the lubrication layer between the grid and the filter paper

We used a confocal light microscope to measure the distance between the nearest fibers of the filter paper and the holey-carbon film on the EM grid during blotting. This distance corresponds to the thickness of the initially continuous lubrication layer that exists between the grid and the wetted filter paper, as shown in Figure 1B. To do this, we constructed a small, variable-pressure clamp to replicate the conditions with which the filter paper is pressed against the grid when inside the Vitrobot.

The pressure exerted by filter paper on the Vitrobot paddles was first measured by replacing the grid with a flat, force-sensitive resistor in a voltage divider and running the regular blotting protocol on the Vitrobot. The measured voltages were calibrated to pressures exerted by known weights. With the Vitrobot at the middle of its force range (blotforce = 0), 20kPa was measured on the resistor. This value varied by less than 2kPa over the available range of the Vitrobot, so 20kPa was chosen as the applied pressure for our blotting experiments, which were performed outside the Vitrobot.

The fibers of pieces of the commonly used #1 Whatman filter paper were stained using 20µL of 100µM Rhodamine B and allowed to dry. A plasma-treated holey-carbon grid (Quantifoil Micro Tools, Jena, Germany) was placed onto a glass coverslip with the carbon film facing up, followed by 20µL of deionized water. The excess of water was used to prevent the sample from drying in the brief moment between deposition and image capture. The stained piece of filter paper was then laid onto the wet grid, an additional coverslip was laid onto the filter paper, and pressure was applied by clamping the sandwiched sample using springs adjusted to apply 20kPa.

Spinning disk confocal microscopy was used to measure the distance between the carbon film of the grid and the stained fibers of the filter paper. The samples were illuminated with a 560nm laser and a vertical stack of images was captured using a 40x NA 0.65 objective. Over each 50µm square of the grid, the distance between the carbon film and the nearest fiber of the filter paper was measured by comparing the z location of the frame with the holes of the carbon film in focus and the z location with the nearest edge of the nearest fiber in focus.

### High-speed interferometry of the formation of thin samples

Reflection interference contrast microscopy (RICM) (10) was used to measure variations in the thickness of water that remains on different areas of a hydrophilic substrate during and after blotting. The EM grid was replaced by a 3mm diameter glass coverslip for these experiments because of practical limitations of the image quality when imaging through a grid.

The coverslips were washed with 100% ethanol and dried with nitrogen, followed by treatment with an oxygen plasma using the March Plasmod System (Nordson March, Concord, CA) for 30s immediately prior to use, which produced a hydrophilic surface. The advancing contact angle of less than 2 degrees on treated bare glass was confirmed to be within experimental error of that on a glow-discharge treated carbon film. The contact angles were measured by depositing a 0.5 µL droplet of deionized water on each type of substrate, measuring the area of the droplet after it had spread, and modeling the volume of water as forming a spherical cap.

Coherent illumination was created on a widefield inverted microscope by passing light emitted from a mercury arc lamp through a pinhole. The coherent light was passed through a 560 nm narrow bandpass filter to a 50/50 beam splitter, which produced an interference pattern on the camera that was dependent on reflection from the sample surface. High-speed images were captured using a Photron 1024PCI (Photron, Tokyo, Japan) camera.

The 3 mm-diameter glass coverslips were placed either onto a filter paper with a hole large enough to image through, or directly onto a glass slide. In either case, 3 µL of 1× phosphate-buffered saline or water was pipetted onto the top of the 3 mm coverslip, to accurately simulate the practice done when blotting EM grids. Blotting was performed by attaching a piece of filter paper to a brass weight, sized to apply 20kPa, and placing it onto the coverslip to produce the desired pressure over the coverslip.

## RESULTS

### Variable electron transmission and ice thickness

The commonly observed variability of ice thickness on an EM grid prepared using blotting can be easily observed at low magnification in a TEM, where it produces a markedly variable degree of electron transmission. The example shown in Figure 2A and Figure 2B, referred to as a “grid atlas”, is not exceptional. Some regions, typically encompassing many grid squares, are completely opaque. The image occasionally is especially bright in other grid squares, indicating the support film broke and is missing from those places. In addition, there are variable degrees of image brightness and contrast in the remaining, quite transparent areas of the grid, corresponding to areas of variable ice thicknesses. In some areas, the grid squares are transparent right to the edges of the grid bars, while in other areas the transparent parts of the squares are confined to a greater or lesser degree, sometimes even being circular. This indicates that the ice is thin in the middle of the square, but thicker ice remains adjacent to the grid bars.

**Figure 2.**
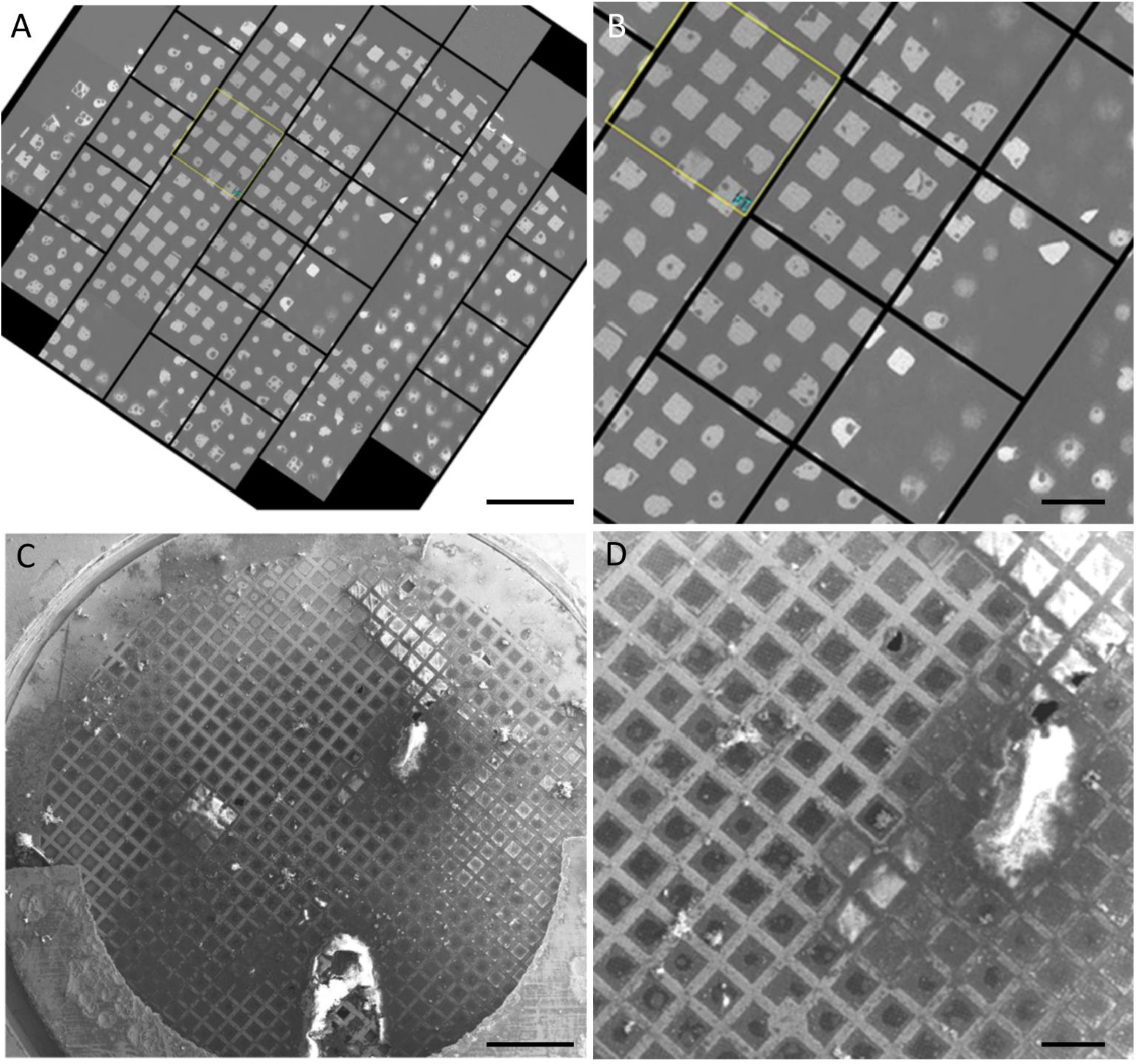
Example of variations in ice thickness over a grid. (A) Cryo-EM “grid atlas” micrograph of an aqueous film prepared using blotting, showing regions of high and low electron transmission. Scale bar=400µm (B) A region of interest from the grid atlas, shown at higher magnification. Scale bar=100µm (C) Cryo-SEM image of the same specimen, with several regions of thick ice, corresponding to local regions of low electron transmission. Scale bar=400µm (D) The same region of interest from the cryo-SEM image. Scale bar=100µm

Cryo-SEM images of the front (Figure 2C,D) and back (Figure 3) surfaces of this same grid confirm, as expected, that areas of low electron transmission correspond to areas with greater ice thickness. In addition, these SEM images also provide information about the side of the grid on which the ice is located, which might not have been guessed from just a projection of the total thickness like that shown in Figure 2A.

**Figure 3.**
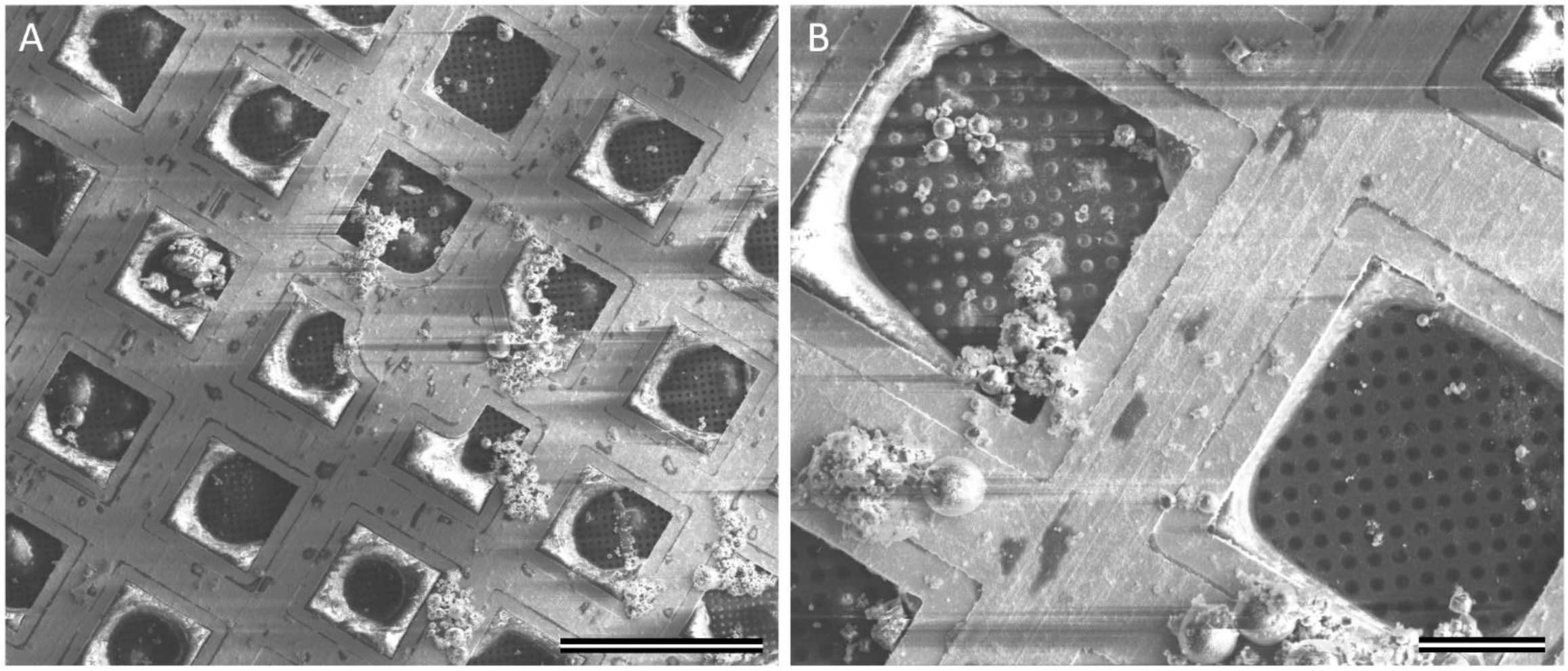
Cryo-SEM image of the back side of a grid, showing thick ice on the perimeter of the grid squares. (A) Scale bar =100µm (B) Scale bar =20µm

The SEM image of the top (or holey-carbon) side of the grid (Figure 2C,D) shows a few irregular, thick regions (“puddles”) of frozen sample, which cover even the metal grid bars and correspond in size and shape to some of the opaque areas in the TEM “grid atlas”, Figure 2A. The ice over most of the rest of the top surface of the grid appears to be relatively thin, as is evidenced by the absence of strong charging effects in the image. Some of the carbon-film squares appear to be covered by thicker ice. This thick ice seems to show charging effects, which manifest as bright white regions covering either the entire square or just regions closest to the grid bars, but not the grid bars themselves.

When viewed from the top (Figure 2D), many of the grid squares have regions of variable width, adjacent to the grid bars, in which the SEM contrast is slightly different than it is in the center. The TEM “grid atlas” in Figure 2A shows exactly the same regions of variable width around the perimeter of otherwise transparent squares, and, in this case, it is clear that the ice is much thicker near to the grid bars than it is in the center of the square. The SEM image of the back side of the grid (Figure 3) shows that the ice adjacent to the grid bars is actually on the back side. This suggests that some of the sample can make its way to the back side of the grid before or during blotting. When that happens, some of the sample can apparently be retained after blotting, where it simultaneously wets the grid bars and – to a variable extent – the region of the carbon film that is closest to the grid bars.

### Initial lubrication layer thickness

We next used confocal light microscopy to measure the proximity of the filter-paper fibers to the carbon film, as described in the Methods. Excess solution was used in this case to prevent drying while capturing the confocal image stacks. We found that a layer of aqueous solution thin enough for use in cryo-EM is not formed simply upon contact between the grid and filter paper. Instead, at least in the visible spaces between copper grid bars, the distance between the nearest fiber and the holey carbon film was typically multiple micrometers (Figures 4A, B). Of the 21 grid squares imaged, only two displayed fibers within a micrometer of the carbon film. Because the first fibers to touch the grid tend to hold the filter paper away from the carbon film under typical blotting pressures, the lubrication layer upon contact between the grid and filter paper is significantly thicker than what is required for cryo-EM. This suggests that some additional process must occur that thins the lubrication layer after full contact is made between the grid and filter paper.

**Figure 4.**
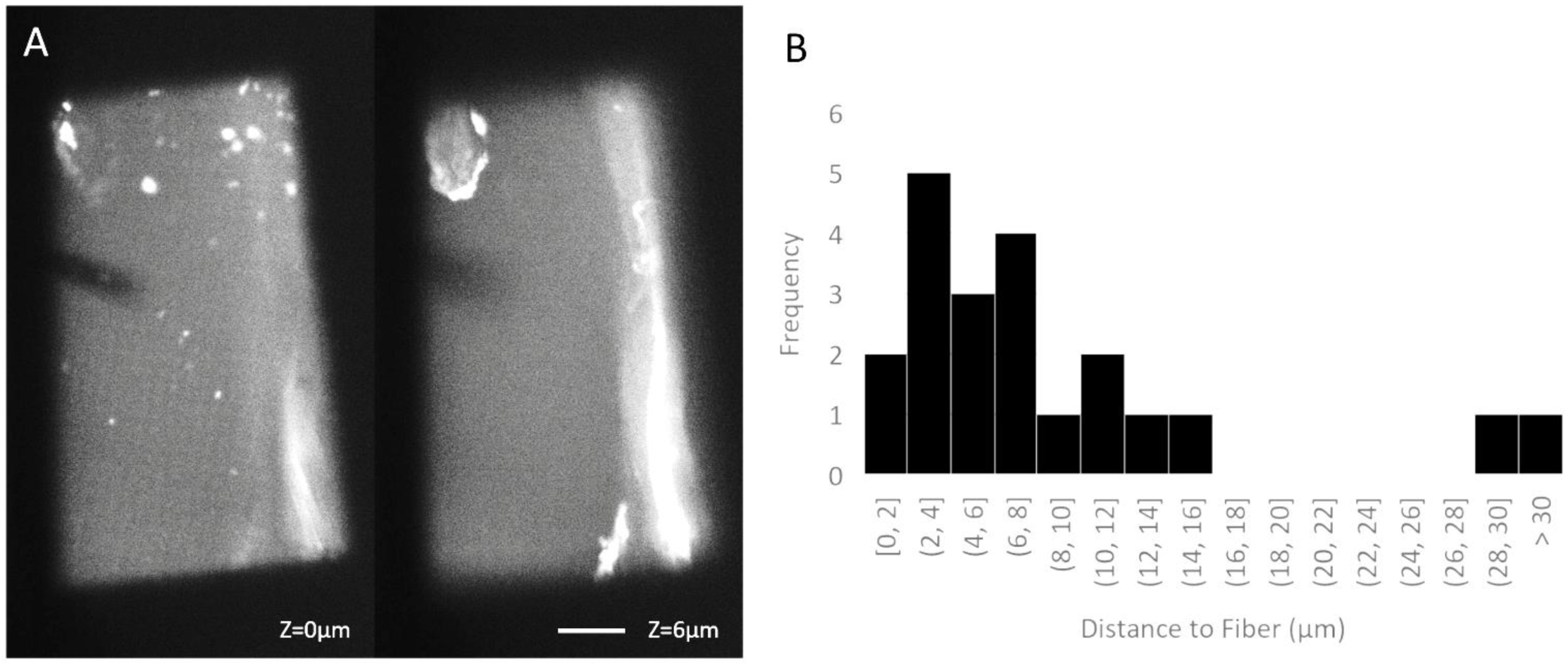
Thickness of the lubrication layer between the carbon grid and filter paper fibers is measured using confocal fluorescence microscopy. (A) Typical images collected showing the holey carbon film (left) and nearest fiber (right) in focus. Scale bar=10µm, DOF=1µm (B) Distribution of distances between carbon film and the nearest filter paper fiber within the 50µm grid square, measured with 20kPa applied. The nearest fiber is typically at a distance of multiple microns above the carbon film.

### Propagation of air into the interface between grid and filter paper

Identifying the mechanism of further thinning required that the process be observed in real time, using typical volumes of aqueous sample. To do that, we used high-speed reflection interference contrast microscopy (RICM) (Figure 5A). We recorded changes in the aqueous-layer thickness during blotting of a 3-microliter sample deposited onto a hydrophilic 3-mm diameter glass coverslip, used as a proxy for an EM grid. After initial contact is made between the coverslip and the filter paper, there is a continuous lubrication layer, as before, and no interference fringes are observed (Figure 5B). However, this continuous lubrication layer lasts for only tens of milliseconds. Then, we see multiple fingers of air propagating in from the edges of the sample towards the center (Figures 5C, D), replacing the liquid that is absorbed into the filter paper. The propagating front of air often pauses momentarily at fibers that are close to the coverslip, but then it quickly jumps tens or hundreds of micrometers to the next fiber (Supplemental Movie S1). The velocity of the advancing meniscus frequently reaches 0.1 m/s during such jumps. After the propagating fingers of air have first entered a region, they continue to broaden laterally as water continues to be absorbed into the filter paper.

**Figure 5.**
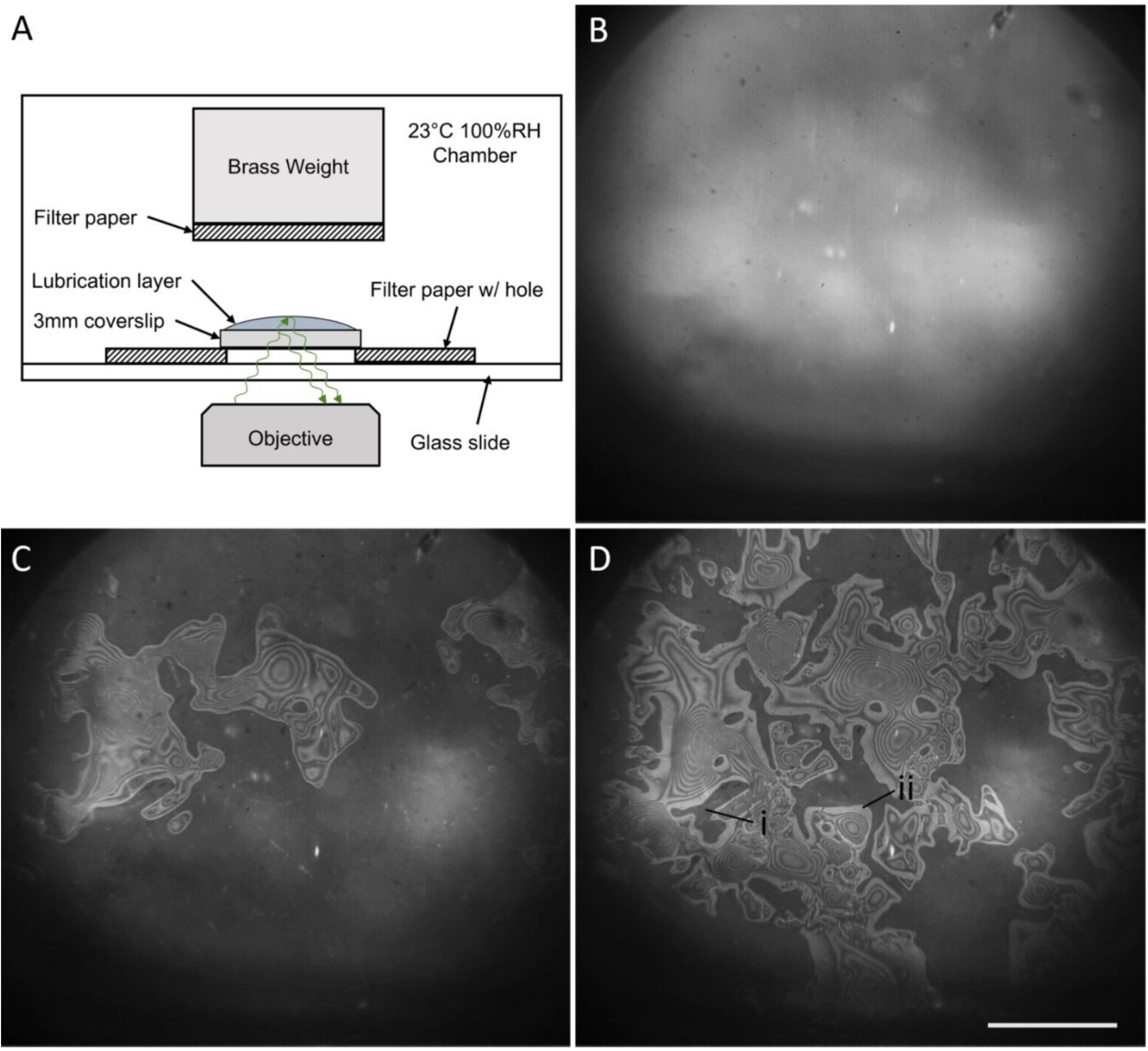
High-speed imaging of water removal during blotting (A) Schematic of the experimental set-up to produce optical interference bands, indicating the height of the lubrication layer. (B) Continuous lubrication exists immediately after contact with the filter paper. (C) Fingers of air begin to propagate from the edges of the grid as capillary pressure draws water deeper into the filter paper. (D) Air continues to propagate. Aqueous solution above region ‘i’ remains continuous with the wetted filter paper, while region ‘ii’ is possibly dewetted. Scale bar =500µm

In certain regions, such as the one labeled “i” in Figure 5D, no air enters the region until the filter paper is removed. These regions are dark in the image because there is no nearby air-water interface to reflect light. We believe that a fiber must be very close to the coverslip in such regions.

On the other hand, there are especially bright regions at the frontier (perimeter) of most of the intruding fingers of air, such as the one labeled “ii” in Figure 5D, These bright bands tend to be wider than the usual interference fringes, and their widths generally increase as the fingers themselves expand in area. We infer that the water thickness within the bright perimeter must be less than 100 nm, since the thickness increases by only 210 nm when going from one bright interference fringe to the next (and a thickness of 105 nm would result in a dark fringe). Such very thin, quite possibly even dewetted regions are expected to manifest as being the brightest ones because of the significant reflection resulting from the mismatch of the index of refraction between glass and air.

Finally, we observe that expansion of the fingers of air leaves behind caps/puddles of water over a majority of the substrate. The thickness in these regions is often one or more micrometers, as can be inferred by counting the interference fringes from the perimeter to the center of a cap. These caps are surrounded by the very thin margins that were described above. Because the water, if any, in these margins is so thin, we suggest that they must impede the flow (removal) of the thicker water from the caps into the filter paper. While the water remaining in the caps is much thinner than the average gap between coverslip and filter paper, what remains is still generally too thick for such areas to be used for high-resolution cryo-EM. As we explain later, in the Discussion, the comparatively large fraction of the area where this is the case implies that a coverslip may not be a perfect proxy for an EM grid.

### Relaxation of the air-water interface after removal of filter paper

As Figure 6 shows, the stable pattern of interference fringes that is established, when blotting has virtually ceased, again undergoes a sudden change when the filter paper is removed. As is expected, a major reorganization of the remaining water occurs when the constraint (i.e. boundary condition) to simultaneously wet the substrate and nearby fibers of the filter paper is removed. Indeed, the rupture of aqueous bridges between the substrate and the wet filter paper might be imagined to be a locally violent event.

**Figure 6.**
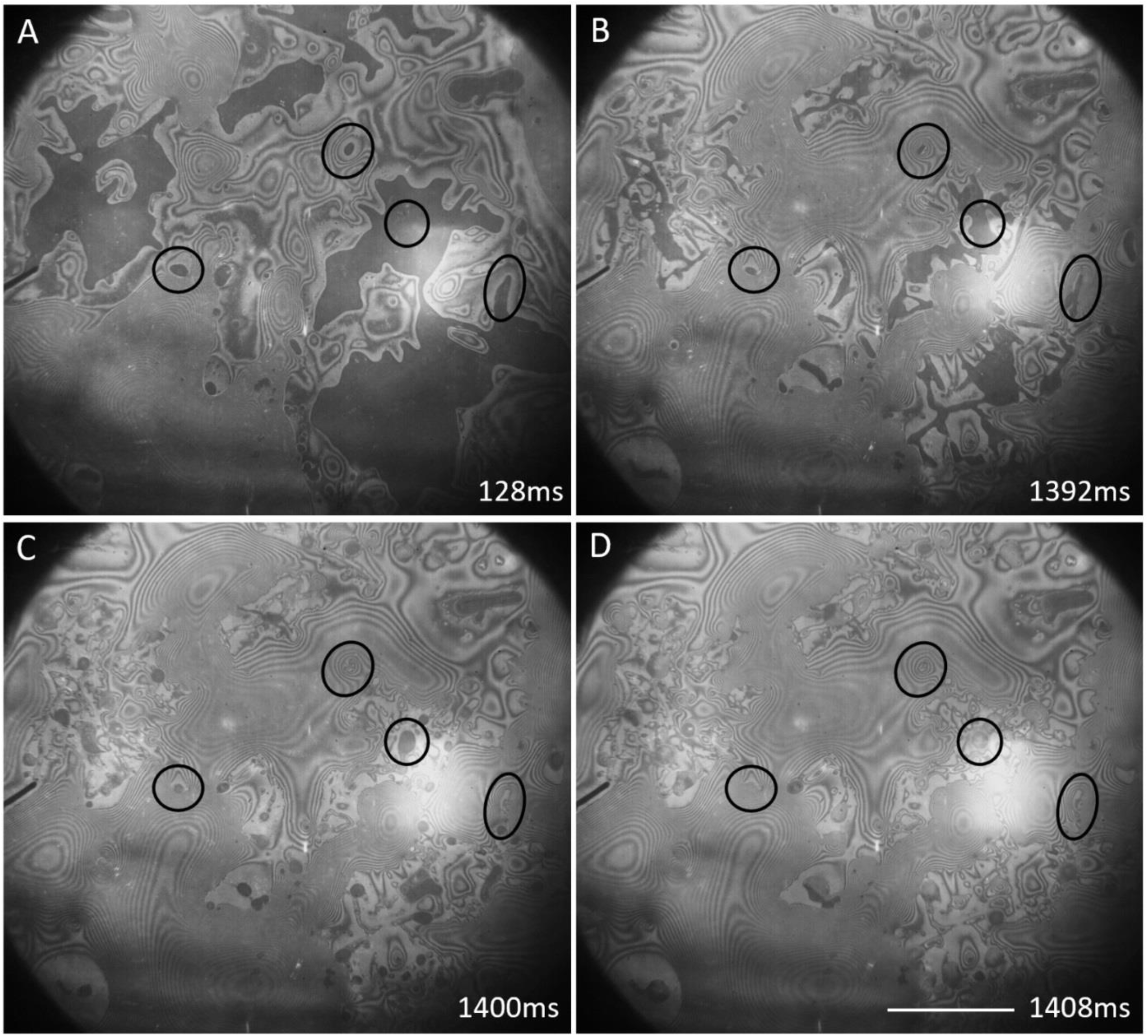
Local relaxation of the air-water interface occurs after the filter paper is removed. (A) Filter paper is still in contact with the substrate. Circled regions indicate examples of fibers near the grid that are constraining a continuous lubrication layer. (B) The filter paper is still in contact with the substrate and additional liquid has drained. (C) Immediately after the filter paper is removed, surfaces that were previously bounded by a fiber collapse to flatter geometries. (D) The surface continues to collapse to a flatter geometry, but apparently dewetted regions on the substrate prevent complete collapse. Scale bar = 500µm

Among the changes that are noteworthy, we point out that the distribution of bright (thus very thin, perhaps even dewetted) regions, which are located at the very perimeter of fingers of air when the filter paper is still in place (Figures 6B), remains relatively undisturbed in some areas but changes considerably in other areas after the filter paper is retracted (Figure 6C and 6D). In addition, the abundance of interference fringes that remains, after the filter paper is retracted, makes it clear that much of the water still remains on the coverslip rather than being pulled off with the filter paper.

It might be considered to be surprising that the remaining water does not merge into a single “spherical cap”, as would be the case if the same total volume were pipetted directly onto a hydrophilic surface. Instead, much of the remaining water is organized into a few, relatively large, irregularly shaped caps, reminiscent of what is often seen in low-magnification TEM images (i.e. the grid atlas). At the same time other, quite thin, regions show a nearly chaotic pattern of interference fringes. These thinner regions co-exist with the thicker puddles. Local relaxation to lower energy topology is pronounced, however, in regions that previously surrounded a fiber in close proximity with the coverslip, such as those circled in Figures 6C and 6D. The continuous layer of water under such fibers collapses to a flatter dome within tens of milliseconds (Supplemental Movie S2). Dewetted regions, which often occur adjacent to fibers, appear in some places to serve as boundaries, preventing the relaxation of the surrounding topology after removal of the filter paper.

## DISCUSSION

### Behavior observed with a coverslip partially captures the results seen after blotting EM grids

We believe that what is observed with a coverslip is instructive about much of what happens when blotting EM grids. It is likely, for example, that fingers of air intrude at random into the well-lubricated gap between filter paper and an EM grid, just as they do for a coverslip. In addition, it is likely that poor control of the thickness of water that remains is inherent to the spatially random way in which fibers of the filter paper must approach the support film on an EM grid.

Nevertheless, the water remaining over most of the area of the coverslip, which we observe using light microscopy, is often thicker than what is usually seen on EM grids during transmission EM. The interference fringes shown in Figures 5 and 6 represent contours of constant thickness that differ by ∼210 nm from one bright fringe to the next (or equivalently, from one dark fringe to the next). Since puddles of water on our blotted coverslips often contain 10 or more sets of fringes, the thickness of water remaining over much of the coverslip can be up to 2 µm, which is effectively opaque to 300 keV electrons. What is more, there are only a few areas of a coverslip where the water is as thin as 100 nm or less, whereas such areas often occur more extensively on vitrified EM grids. These differences imply that the outcome after blotting a coverslip, used as a proxy for an EM grid, does not tell the full story of what happens during blotting EM grids.

We recognize that there are a number of physical differences between a 3-mm glass coverslip and a 3-mm EM grid, even if the two surfaces have been made equally hydrophilic, as was the case here. For example, the surface topology of an EM grid is not as uniformly flat as is that of a glass coverslip. Instead, the holey carbon film on an EM grid may sag into the open space between grid bars, something that is often noted to be the case for Quantifoil grids. As a result, the fibers of filter paper are likely to contact mainly the metal bars of an EM grid, but not the holey carbon film. Another significant difference is that the thin carbon film is a much more pliable substrate than is a rigid coverslip. As a result, suction of water into the filter paper may draw the carbon-film substrate into more conformal contact with the irregular filter paper. Finally, the back side of the carbon film seems to always become wetted during blotting, if not before, a feature that is not captured when using a coverslip as a proxy. It may be that one or more of these differences can account for the greater retention of water on a coverslip.

### Modeling how water is removed from EM grids by blotting

As indicated in Figure 1B, blotting is expected to initially produce a continuous “lubrication layer” of water that is in contact with the entire surface of the EM grid and extends into the filter paper. This is expected to happen even if the plane of the filter paper is initially inclined by a small angle relative to the plane of the grid, as it is by design in the ThermoFisher Vitrobot instrument but not in the Leica EM GP2 Automatic Plunge Freezer. It is currently unknown, however, whether tilting the plane of the filter paper relative to that of the grid subsequently results in microscale fluid behavior that is qualitatively different from what we observed here, with no tilt. It is true that a small tilt might be expected to cause air fingers to consistently penetrate the lubrication layer at the more open end of the gap between grid and filter paper. Furthermore, if the slope is large enough, fibers from the filter paper would no longer impose constraints on removal of water from the areas of the grid where the gap is large enough. The extent to which an initial slope matters, however, may depend upon the extent to which the filter paper is initially drawn into flat contact with the EM grid by capillary action, especially when the slope of the filter paper is quite shallow. In any event, we are not aware that the cryo-EM community has found that existing implementations for blotting grids produce qualitatively different final results, suggesting the process may be insensitive to a small angle of inclination of the filter paper.

How much sample is subsequently removed may reasonably be expected to depend upon the blotting pressure exerted between the grid and the filter paper. We also expect that it will depend upon the capillary pressure exerted by the pores of the filter paper relative to the capillary pressure exerted by the gap between the filter paper and the hydrophilic substrate. What is difficult to predict, however, is whether the filter paper and substrate remain well-lubricated as liquid is wicked into the filter paper, thus causing the gap between them to become smaller, or whether the size of the gap remains almost unchanged and air replaces liquid as the latter is drawn into the filter paper. In practice, we observe mainly the latter under the preload pressure exerted by the Vitrobot. Much of the lubrication layer is displaced by air that penetrates from the perimeter, but some liquid is permanently retained at points where fibers of the filter paper come very close to, or touch the hydrophilic substrate.

In order to understand the rapidly evolving, irregular pattern of water that results during blotting, an example of which is shown in Figure 5, we first consider an area at the edge of a grid, where the lubrication layer is open to ambient pressure. Figure 7 shows a cartoon representing a cross section of the essential elements. The dashed line in this cartoon denotes a control volume that includes the liquid meniscus between a fiber and the grid and another between two adjacent fibers. This control volume is treated as a small, open channel with different values of the capillary pressure at either end. The two values of pressure are themselves estimated using a variation of the Young-Laplace equation, as follows:

**Figure 7.**
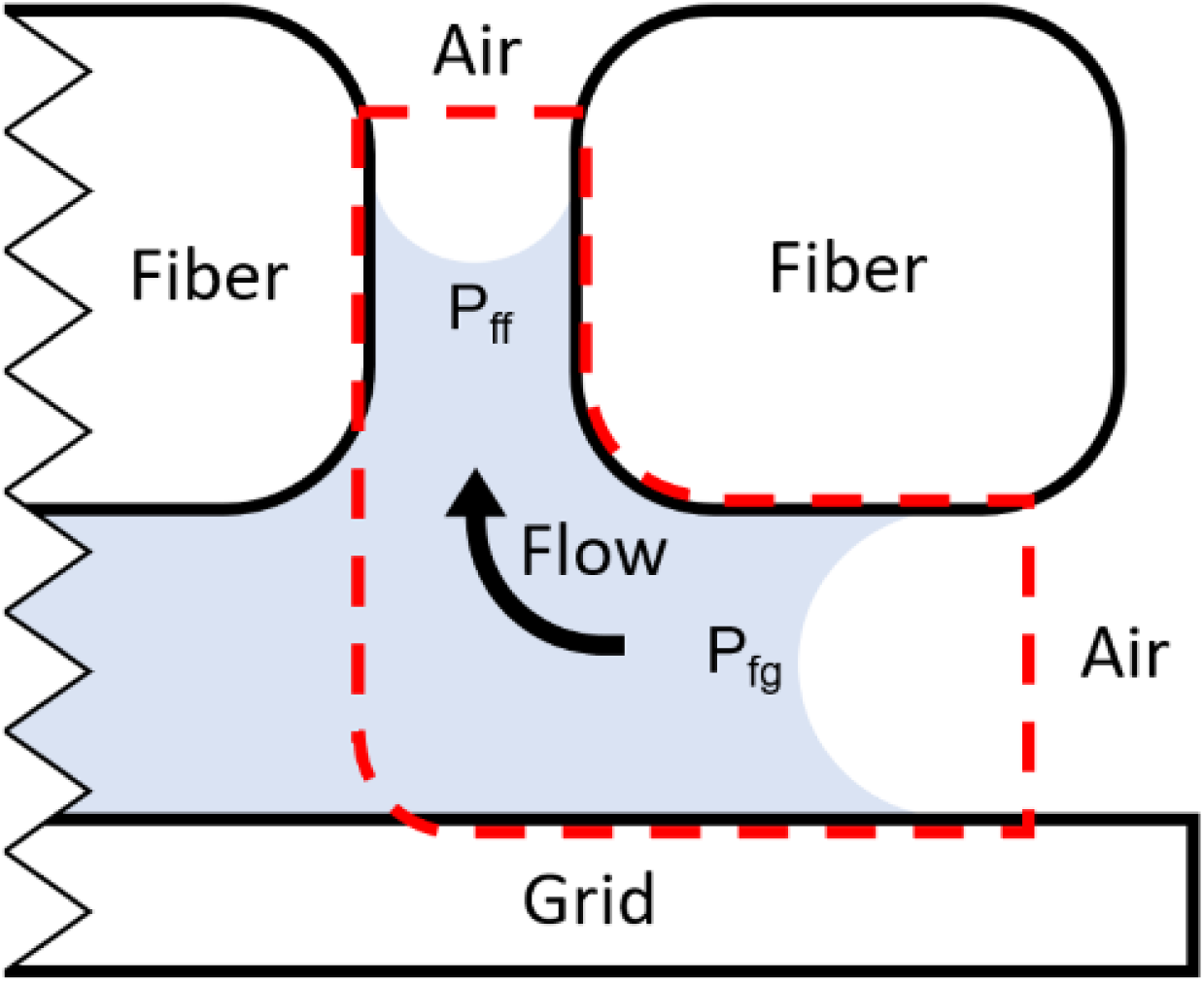
Schematic cross section of the grid – filter paper interface at the edge of the grid. Imbalance in the capillary pressures imposed between two fibers and between the grid and a fiber results in the propagation of air from the edges of the grid towards the center and upwards into the filter paper.

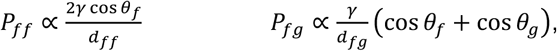

where *P*_*ff*_ is the capillary pressure between two adjacent fibers, *P*_*fg*_ is the capillary pressure between a fiber and the grid, *γ* is the surface tension of the liquid, *θ*_*f*_ is the fiber contact angle, *θ*_*g*_ is the grid contact angle, *d*_*ff*_ is fiber-fiber distance, and *d*_*fg*_ is fiber-grid distance.

The distribution of smallest distances between the holey carbon film, spanning a 50 µm square of the grid, and the nearest fiber in the filter paper (Figure 4) suggests that the equivalent capillary widths between the substrate and filter paper fibers will vary locally from less than a micrometer to tens of micrometers. This distribution overlaps with the nominal pore sizes of filter paper, specified by vendors (e.g. Sigma Aldrich) to be ∼7 µm for Whatman grade 595, the standard filter paper used with a Vitrobot and ∼11 µm for Whatman grade 1, used by some groups for blotting grids and also here, for our experiments. Because of this overlap of fiber-grid and fiber-fiber distances, it is inevitable that some regions of the initial, continuous lubrication layer will be bounded by small fiber-fiber distances and large fiber-grid distances, while other regions are bounded by large fiber-fiber distances and small fiber-grid distances. Variation in the distribution of fiber-fiber distances in different grades of filter paper is not investigated here, but it may also result in changes in the geometry and coverage of the air fingers over the grid surface.

The capillary pressure of pores within filter paper is thus expected to be stronger than the capillary pressure between fiber and grid over many areas of the face of an EM grid. We propose that this difference in capillary pressure explains why water is sucked into the filter paper, being replaced by air that intrudes from sites along the perimeter of the grid. Fingers of air then continue to intrude more deeply from those sites into the gap, finding the route of least resistance for ambient pressure to continue pushing water into the filter paper.

In areas where the gap between a fiber and the grid surface is, on the other hand, smaller than the pore size of the filter paper, the capillary pressure within the gap will be stronger than it is within the pores of the filter paper. We reason that liquid in such places will remain “pinned” between grid-surface and fiber at those spots. According to the data shown in Figure 5, intruding fingers of air quickly circumnavigate around such spots, leaving behind only a thin film of water that continues to wet the substrate wherever, we infer, the thickness of the gap must have been larger than the pore size of the filter paper, roughly speaking.

To summarize, we observe that irregular fingers of air propagate from the edges of the grid, leaving behind both randomly located, thick regions of continuous lubrication and regions where thin aqueous films are now separated by an air gap from the wet filter paper. When this process stops, the geometrical shape of the resulting air-water interface will be a “minimal energy surface”, or at least one for which the energy is a local minimum. The liquid below that interface is thickest near to the fibers that remain in close contact with the substrate and thus effectively “pin” the interface at those locations. The thickness of the film of water remaining everywhere else, i.e. between points of contact between fibers and the substrate, is expected to depend, in the first instance, upon how far apart the respective, well-lubricated fibers are from one another, and upon the unique spatial distribution of those points. As mentioned previously, the interference fringes seen in RICM images, such as Figure 5 and Figure 6, serve as constant-thickness contours, with the thickness of water changing by 210 nm from one bright band to the next.

In places where a relatively large area is devoid of fiber-contact points, the curvature of the minimal-energy surface is expected to be small. Since the capillary pressure in such areas is expected to be correspondingly small, liquid is free to be drawn into the filter paper, leaving only minute volumes on the substrate. Such areas are thus at risk of dewetting, and places where that may have happened are seen in Figure 5 and Figure 6. If amphiphilic surfactants are adsorbed at the air-water interface, however, such thin regions can be stabilized by a disjoining pressure (11, 12), and the risk of dewetting can be reduced.

A point that still needs to be addressed further is the fact that the thickness of water that remains over much of the area of a 3-mm coverslip, once blotting is complete, is often greater than that which remains over significant portions of most EM grids after vitrification. One possible explanation that we initially considered is that much of the liquid might be wicked away simultaneously with the filter paper being “peeled” from the coverslip. This hypothesis was shown not to be the case; rather, Figure 6 demonstrates that the thickness profile changes only little, if at all, when the filter paper is removed. Evaporation has always been mentioned as something that could thin the sample after blotting an EM grid. As reported in Figure 4 of (13), however, the evaporation rate is less than 5 nm/s at a relative humidity above 90%, which may safely be expected to be the case if the maximum possible relative humidity is used in a Vitrobot or similar environment. Our own experience has also been that evaporation only trivially changed the thickness profile under the conditions that we set up.

Another point that is worth commenting on is the fact that the topographies of the air-water interfaces on the front and back surfaces of vitrified EM grids are very different, as viewed with SEM. Empirically, it is commonly found that the filter papers on both sides of the grid are wetted after blotting, irrespective of whether spherical caps of water are seen on one or both sides before blotting. The process by which wetting both sides of a grid occurs remains an unanswered question. Putting aside the question of how water, applied to the front side only, also gets to the back side of a grid, it is not at all surprising that a microscale meniscus of liquid often wets each grid bar and the adjacent holey carbon film simultaneously on the back side, as is seen in Figure 3. We also note that the distribution of water remaining within a grid square is often biased in one direction, inviting the suggestion that water was drained from the back side in mainly one direction, perhaps by intrusion of air fingers just as is the case on the front side. We further note that the amount of liquid remaining on the back side of a grid can vary quite considerably from one grid square to the next, as is the case on the front side. It thus seems plausible that similar phenomena to those demonstrated during blotting a coverslip may occur simultaneously on the front and back sides of a grid.

### Changes in water distribution occur only very slowly after removing the filter paper at high humidity

The early literature spoke of the importance of pausing for a number of seconds after removing the filter paper, to allow the remaining thin films to further “drain” (13, 14). In the absence of evaporation, as those discussions had pointed out, van der Waals forces can be responsible for further thinning (i.e. draining) in some areas, with a corresponding increase in thickness occurring in other areas. In other words, films of a pure liquid, when they become too thin, are unstable with respect to forming an asymmetric distribution of mass. While this instability can potentially lead to liquid-film rupture, or to dewetting of solid surfaces, the presence of a surfactant on the air-water interface can generate a disjoining pressure (11) that opposes such events. While no surfactant was intentionally added in our experiments, neither were steps taken to keep air-water interfaces perfectly clean. The latter will also be true under conditions when making EM grids of biological macromolecules in which no surfactant has been added. Furthermore, as mentioned above, the protein itself can adsorb to and denature at the air-water interface. When that happens, the adsorbed material might itself generate a disjoining pressure that opposes rupture and subsequent dewetting of a thin film of water.

It is not expected that residual water would drain downwards under the influence of gravity, as some have speculated in personal communication, since the ∼3 mm capillary-length value for water is much larger than the thickness of the sample remaining after blotting. Another speculation is that remaining sample might drain upwards under the influence of capillary action from the tip of the tweezers. Other ideas have been that acceleration of the grid during plunging, or air shear during plunging, might displace excess sample up to the end that is held in the tweezers. The initial location of the tip of the tweezers is now easily seen in the SEM image shown in Figure 2C, and there is no correlation between where the puddles are found and the orientation of the grid when it was plunged. As a result, we suggest that none of the above-mentioned models determine the spatial distribution of thick ice that is often seen on vitrified EM grids.

### Estimating the peak value of shear stress and its possible consequences

Having observed the motion of fluid during the blotting process, we were interested in estimating the scale of mechanical effects such as shear stress, and whether these might degrade biomolecules. By approximating the propagating fingers of air as long bubbles moving through a capillary, we can gain a rough estimate of the highest shear stress experienced by the adjacent liquid during bubble propagation. We note that the leading edge of the propagating bubble is like that in the Bretherton problem of a long air bubble advancing in an open capillary tube (15). We thus use Bretherton’s estimates for shear stress magnitudes: 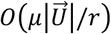 in the region in front of the advancing meniscus, labeled “i” in Figure 8C, and 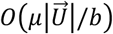 the region labeled “ii” in Figure 8C, where µ is the dynamic viscosity, U is the propagation velocity of the air-water interface, r is half the distance between the upper and lower surface, and b is the thickness of the residual water film remaining on the bottom surface (Figure 8C). Our situation, when blotting with filter paper, is different from the Bretherton problem in that the upper surface in our case is porous, and thus the upper thin film of liquid would be partially drawn further into the filter paper. Although this difference may produce an asymmetric meniscus, rather than the symmetric one shown in Figure 8C, this difference is not expected to have much effect on the shear stress experienced close to the bottom surface.

**Figure 8.**
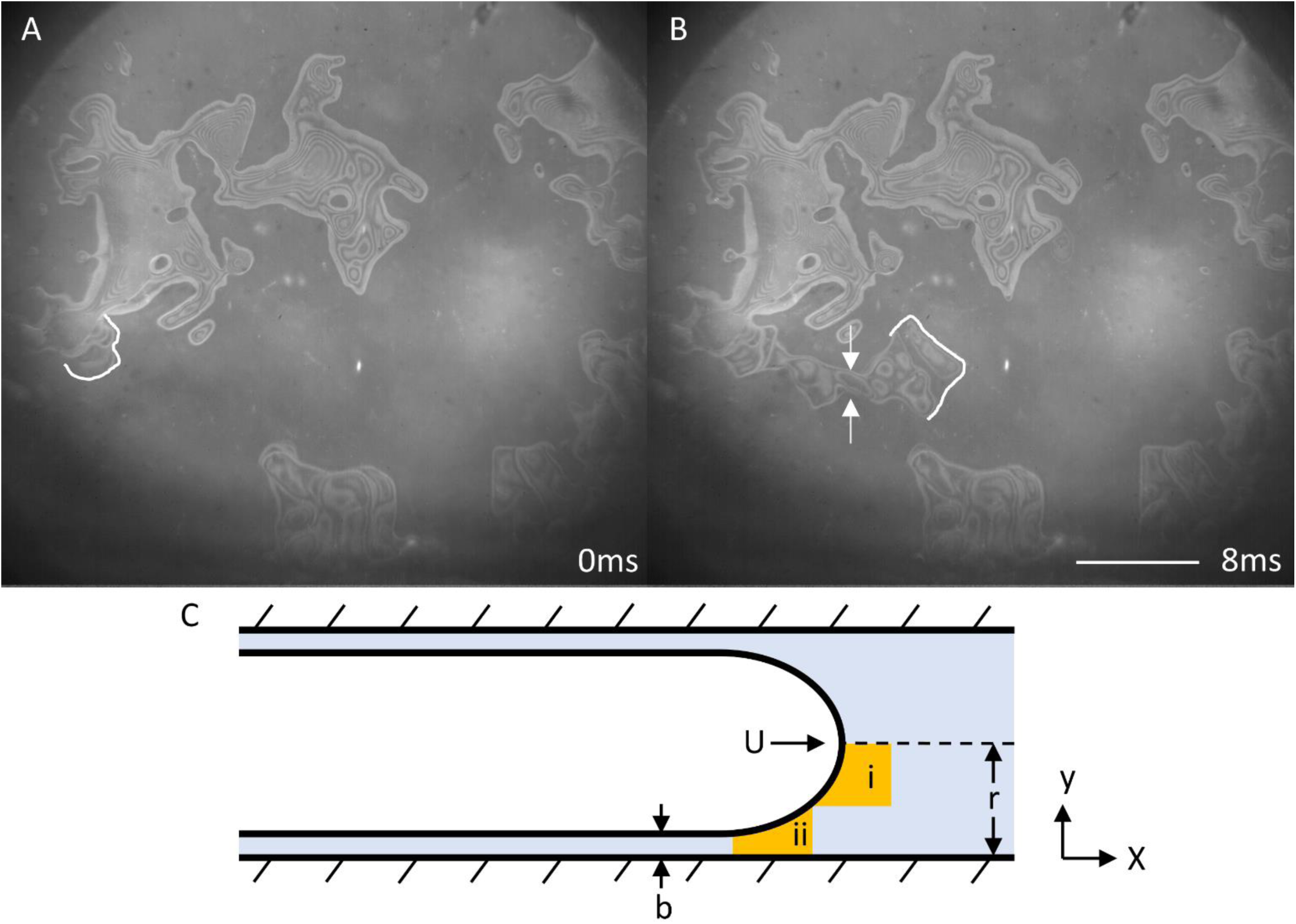
During propagation of the fingers of air, high velocities likely result in significant shear stress in the surrounding liquid. (A) The air-water interface at 0 ms (an arbitrary reference time) is shown in white. (B) After 8 ms, the interface has propagated 750µm through a neck of 75µm diameter, shown between the two arrows. (C) The propagating front of air is modeled as a long bubble in a tube with two regions of interest, i and ii, in which shear stresses are estimated. Scale bar = 500µm

In Figure 8, we examine a typical fast propagation event, with average velocity of 0.1 m/s, fiber-fiber distance of 100 µm, and remaining film thickness of 2 µm. Region i near the center of the pseudo-capillary at any time during the propagation of the interface will experience shear stress around 2 Pa, while region ii, near the substrate may experience 50 Pa. If the remaining film thickness was nearer the final desired thickness for high-resolution cryo-EM, these stresses might be on the order of kilopascals.

While even the largest of these estimated stresses may have no effect on small globular proteins (16), certain larger macromolecules might undergo shear degradation which is not reversed when the shear stress is relaxed. For instance, Von Willebrand Factor, an essential protein in blood clotting, undergoes a dramatic unpacking of its many repeating subunits, which are otherwise loosely clustered together, at 5 Pa shear stress (17), and this activates the protein *in vivo*. However, it remains unclear whether maximum shear stresses estimated here are large enough to result in changes in quaternary structure, at least for some more labile cryo-EM samples. What is well-known, however, is that long filaments such as F-actin, microtubules, and tobacco mosaic virus particles are often seen to be oriented roughly parallel to one another in electron micrographs, something that can only be explained by the flow of buffer that occurs as excess sample is removed. Galkin et al. even reported that F-actin filaments can break during blotting of grids for cryo-EM, and that the degree of order of monomers within F-actin polymers improves when filaments are stretched by flow within the regions of thinnest ice (18).

### Implications for trying to optimize blotting parameters

The commercially available Vitrobot and similar instruments offer access to a number of parameters that the user can adjust in order to optimize the outcome. In addition to the ambient temperature and relative humidity, these parameters may include the blot force, blot time, and pause interval after the filter paper is removed and before the grid is plunged into liquid ethane. Other parameters that users may vary to optimize the result include the volume of sample deposited on grids and the length of time during which the filter paper is equilibrated with the humid environment before blotting occurs. However, there is no standard theory of how these parameters might affect the resulting thickness and uniformity of the final sample.

The results that we present here suggest that the blot pressure at the level of the grid (in the Vitrobot instrument) varies only slightly over the range of blot force provided to the user, and thus we do not expect that parameter to affect the outcome significantly. Furthermore, we observe that the process of removing excess liquid from a 3-mm coverslip is completed in about 1 second or less, after which a static aqueous film remains in equilibrium with the wet filter paper. Since a similar temporal behavior can be expected for a hydrophilic EM grid, it seems that little can be accomplished by blotting for times longer than 2 or 3 seconds. Nevertheless, this time scale might be longer if the sorptivity of the filter paper is reduced significantly, for example by first equilibrating it at high humidity, an issue that we have not investigated here.

Many users choose zero for the pause interval between retraction of the filter paper and plunging the grid into liquid ethane. Even so, there is an inherent (instrumental) pause of about 1 s between retraction of the Vitrobot paddles and plunging. Since we have observed rather little redistribution of remaining water to occur after 1 s, the choice of zero pause seems to be appropriate. An obvious exception, however, would be to add a further pause if one intends to remove water by evaporation.

Although many laboratories may have developed trusted recipes for blotting grids, whether using a Vitrobot or in other ways, our observations suggest that varying many of the available parameters is not likely to change the results obtained. A possible exception, however, is to vary the temperature, which is a parameter that we did not investigate. Furthermore, a significant exception would be to use a lower ambient humidity and a pause time of a second or more in order to allow evaporation to thin the sample, again something that we did not investigate. It is widely recognized, of course, that allowing evaporation to further thin samples may be risky for some proteins, since the ionic strength and perhaps even the pH could change substantially.

The use of scanning electron microscopy has resolved ambiguities about whether regions of thick ice, apparent in the TEM “grid atlas” of the same grid, are all located on the surface to which the liquid sample was first deposited, or whether some also remains on the back side after blotting. In the example shown in Figure 2, rather large regions of electron-opaque ice consist, at least in part, of thick puddles of sample that remain on the front side of the grid. These opaque areas limit the fraction of the grid where one finds open squares that can be used to collect high-resolution data, and it is not uncommon that the reduction in the amount of transparent area can be even more severe than that seen in Figure 2. In addition, thick, even electron-opaque ice often extends out – to variable extents – from the edges of the grid bars toward the centers of otherwise open grid squares. It has sometimes been thought that retention of sample next to the grid bars indicates squares in which high-resolution images can be obtained. However, such a meniscus at the grid bars may be misleading because this ice is located on the back side of the grid. If one uses a continuous support film over holey-carbon, or if there is a sacrificial, denatured-protein skin, like that which is formed by ferritin (19), water on the back side of a grid may not indicate the extent to which sample on the front side still remains embedded in a film of vitrified water.

## SUMMARY AND CONCLUSIONS

Our experiments have observed the rapid intrusion of microscale fingers of air into the fully lubricated gap that initially exists between filter paper and a hydrophilic substrate, shortly after the filter paper is pressed against the substrate. We identify the rapid intrusion of air into this gap as the physical process that leaves behind thin films of sample, even before the filter paper is removed. These observations identify the fact that the boundary conditions created by the filter paper, which govern removal of excess sample during this process, are poorly controlled. This fact is proposed as the explanation for why the resulting ice-thickness values are so variable. Continued work is needed to design and implement alternatives to blotting, such as spraying nanodrops, dip-pen spreading, blotting from the perimeter rather than the face of a grid, or other approaches that may yet be proposed.

## AUTHOR CONTRIBUTIONS

JT,B-GH and RMG recorded SEM images of grids used previously for cryo-EM by B-GH and RMG. DAF and MA originated the concepts for experiments using confocal light microscopy and high-speed reflection interference contrast microscopy. SG contributed to the theoretical analysis of liquid flow in capillary spaces and assisted with experiments. Major portions of the initial draft were written by MA and RMG, with material added by DAF. All authors reviewed and helped to revise the final draft.

## ACKNOWEDGEMENTS

We thank Prof. Stephen Morris for discussions that provided crucial background information. Dr. Mike Vahey participated in early experiments leading up to the work presented here. This work was supported in part by NIH grant P01GM051487 (RMG and BH), and by Chan Zuckerberg Biohub (DAF). Work at the Molecular Foundry was supported by the Office of Science, Office of Basic Energy Science of the U.S. Department of Energy under Contract No. DE-AC02-05CH11231.

